# PFKFB3 protein in adipose tissue contributes to whole body glucose homeostasis

**DOI:** 10.1101/2024.10.17.618949

**Authors:** Beth A. Griesel, Ann Louise Olson

## Abstract

Age-dependent changes in adipose tissue are thought to play a role in development of insulin resistance. A major age-dependent change in adipose tissue is the downregulation of key proteins involved in carbohydrate metabolism. In the current study, we investigate the role of 6-phosphofructo-2-kinase/fructose-2,6-bisphosphatase 3 (PFKFB3) a key governor of the rate of glycolysis in adipocytes via the synthesis of fructose-2,6-bisphosphate that was significantly down-regulated in aged mice. We employed an adipocyte-specific PFKFB3 mouse line to investigate the role of PFKFB3 on adipocyte function. In both aged mice and PFKFB3-knockout mice, we observed an increase in O-glcNAcylated proteins consistent with a shift in glucose metabolism toward the hexosamine biosynthetic pathway. Under chow-fed conditions, PFKFB3 knockout resulted in significantly smaller adipocyte area, but no difference in total fat mass. While glucose tolerance was unchanged under chow conditions, when mice were challenged with a 4 wk high fat feeding, PFKFB3 deletion led to a greater decrease in glucose tolerance as well as a significant increase in macrophage infiltration. These results indicate that perturbation of the glycolytic pathway in adipose tissue has multiple effects of adipocyte biology and may play a significant role in metabolic changes associated with aging. Results of this student support the notion that changes in glucose metabolism in adipose tissue impact whole body metabolism.

## Introduction

Age is one of the major risk factors for the development of Type 2 diabetes, but the mechanisms underlying this risk are unknown. With the steep rise in obesity rates, the development of type 2 diabetes has increased, leading to the notion that adipocyte dysfunction that is observed in both aging and obesity may play a role in disease risk (1). Evidence supports the notion that avoidance of obesity through interventions such as calorie restriction plays an important role in healthy aging (2). This implies that overall expansion of adipose mass or changes in adipose function with aging may contribute to age-dependent disease risk.

Adipose tissue is one of the few tissues that develops postnatally, therefore, increases in adipose number are expect during growth are expected. Following adolescence, the adipocyte number is set and changes in adiposity reflect changes in lipid turnover (3). Long-term changes in lipid turnover measurements in humans revealed that lipid removal decreases as a function of age without a reciprocal adjustment in lipid uptake (4). This imbalance in lipid uptake versus removal leads to increased weight gain with age which may play a role in diabetes risk.

Adipose tissue plays a key role in maintaining metabolic homeostasis and insulin sensitivity by providing a metabolically safe place to store lipid and serving as a source of insulin-sensitizing adipokines such as adiponectin (5, 6). Increased prevalence of obesity and obesity-related diseases as a function of age has increased the need to understand the mechanisms that underlie development and maintenance of healthy adipose depots.

Over the course of the daily transitions between the fed state and the fasted state, adipose tissue undergoes a metabolic transition from a state of lipid storage to a state of lipid release. This transition is orchestrated by changing from high insulin signaling to high catecholamine signaling. Changes in lipid metabolism are accompanied by changes in glucose uptake and metabolism, which may also govern the transition from lipid uptake to lipid release (7–9).

Glucose metabolism plays a key role in the development of adipocytes from progenitor cells (10, 11). We have shown that adipocyte maturation was dependent on the acceleration of glycolysis that was concurrent with the upregulation of 6-phosphofructo-2-kinase/fructose-2,6-bisphosphatase 3 (PFKFB3), a PFK2 isoform that is a key regulator of glycolysis. PFKFB3 catalyzes the production of fructose-2,6-bisphosphate (F-2,6-BP) to allosterically activate phosphofructokinase 1 (PFK1), the committed rate limiting step for anaerobic glycolysis. PFKFB3 is a bifunctional enzyme with kinase activity and phosphatase activity within the same peptide chain. In the case of PFKFB3, the kinase activity is ∼700 more active than the phosphatase, thus favoring F-2,6-BP production (12). PFKFB3 is expressed at high levels in adipose tissue compared to other tissues (13, 14). PFKFB3 protein expression is reduced in adipose tissue of fasting animals and rapidly increased after 4 hours of refeeding (10).

In the current manuscript, we show that PFKFB3 is significantly downregulated in adult mice at least as early as 7 mos of age. To study the impact of decreased PFKFB3 expression specifically in adipocytes, we developed an adipocyte-specific knockout of PFKFB3 mouse model. Our results revealed that PFKFB3 deficient adipocytes were smaller despite normal fat content. Both PFKFB3-deficient you mice and non-transgenic aged mice had increased O-GlcNAcylated proteins. When challenged with a short-term high fat diet, adipose-specific knockout of PFKFB3 result in impaired glycemic control and increased macrophage infiltration. Taken together, these data suggest that PFKFB3 activity in adipocytes may be required for optimal storage and handling of nutrients, which is perturbed with aging.

## Materials and Methods

### Animals and diets

Transgenic animals used for these experiments were male and female C57Bl/6 mice bred to carry an adipose-specific conditional knockout of the gene encoding PFKFB3. This was carried out by crossing mice carrying a homozygous floxed pfkfb3 allele with B6.FVB-Tg(Adipoq-cre)1Evdr/J mice from Jackson lab (RRID:IMSR_JAX:028020). All procedures using animals were approved by the Institutional Animal Care and Use Committee at the University of Oklahoma Health Sciences Center.

Mice were kept in a temperature-controlled room with a 12-hour light/dark cycle. From weaning to twelve weeks after birth, the transgenic and floxed control mice were group housed and ad libitum fed either chow diet (RD) (13% kcal from fat, 5053 Rodent Diet, LabDiet) for lean mice. Some mice were then switched to an obesogenic high fat diet (HFD) (60% kcal from fat; D12492, Research Diets Inc., New Brunswick, NJ) for 4 wks to induce obesity. All mice were given free access to water.

For aging studies, C57Bl/6 wild type male mice were group housed and ad libitum fed RD for 2 mos, 7, mos, or 20 mos of age. All mice were given free access to water.

### Western blot analysis

Total tissue extracts from adipose tissue and whole cell extracts from RAW 264.7 (American Tissue Culture Collection, Manassas, VA; RRID:CVCL_0493) were prepared in lysis buffer containing 20mM HEPES, 1% NP40, 2mM EDTA, 10mM sodium fluoride, 10mM sodium pyrophosphate, 1mM sodium orthovanadate, 1mM molybdate, protease inhibitor cocktail (complete Mini EDTA-free Protease Inhibitor Cocktail, Roche), 1 mM PMSF and 50 mM DTT. Protein concentrations were determined by using a Coomassie Plus (Bradford) Assay Kit (Pierce). Lysates were fractionated using SDS-PAGE, and proteins transferred to Immobilon-FL polyvinylidene fluoride membrane (EMD Millipore Corporation, Billerica, MA, US.) Membranes were stained with anti-GLUT4 antibody (ab33780, ABCAM; RRID:AB_2191441), anti-adiponectin antibody (a gift from Philipp Scherer, UT Southwestern Medical Center, Dallas TX), anti-PFKFB3 (ab181861, ABCAM;RRID:AB_3095816), anti-F4/80 antibody (ab300421, ABCAM; RRID:AB_2936298), anti-α/β-tubulin antibody (2148, Cell Signaling; RRID:AB_2288042), anti-O-GlcNAc (MA1-072,Thermo Fisher Scientific; RRID:AB_326364), or anti-O-GlcNAc transferase (Sigma-Aldrich, 06264; RRID:AB_532313) and visualized with appropriate secondary antibodies conjugated with AlexaFluor 680 (Invitrogen). Rabbit anti-lipoic acid antibody for pyruvate dehydrogenase subunit E2 (PDH-E2) and alpha-ketoglutarate dehydrogenase subunit E2 (KGDH-E2) immunolabelling was provided by Kenneth Humphries (Oklahoma Medical Research Foundation) (15) Fluorescence was quantified with a Li-Cor Odyssey imager (Li-Cor Biosciences, USA). Protein expression data were compared to tubulin as a loading control. Membranes were visualized using the appropriate AlexaFluor 680-conjugated secondary antibody and quantified using the Odyssey imaging system (Li-Cor Biosciences).

### Plasma adiponectin

Plasma adiponectin was measured in random fed mice using the mouse adiponectin ELISA kit (Crystal Chem, 80569).

### Immunofluorescence and Adipocyte area

Epididymal fat pads harvested from all groups of chow fed mice at 3 mos of age or at 4 mos of age after 4 weeks of high fat diet and were fixed in 10% neutral buffered formalin, embedded in paraffin and cut into 5 µm sections. Macrophage infiltrates were stained with anti-F480 antibody (ab300421, ABCAM; RRID:AB_2936298) and visualized with a secondary antibody conjugated with AlexaFluor 647. Three random fields from 3 independent mice per group were collected. Images were collected using a Nikon Eclipse Ti fluorescent microscope and analyzed with Nikon Elements software.

Adipocyte area was measured in hematoxylin and eosin-stained paraffin sections of perigondal adipose tissue. Images were collected using a Nikon Eclipse Ti fluorescent microscope and analyzed with Nikon Elements software. The largest 10 unbroken cells per section were counted, as these were thought to be more likely to represent the true diameter of the cells. Four areas from three different mice were measured for a total of 120 cells.

### Glucose tolerance test

After a 6 h fast, mice were given 2 mg/kg i.p. glucose and tail vein blood glucose was measured using a TRUETrack glucometer (Walgreen’s) that was sche. Blood glucose was measured at 0.25, 0.5, 1, 1.5, 2, and 3 hours after injection. Area under the curve was measured using GraphPad Prism.

### Triglyceride and glycogen measurements

Lipids were extracted from liver using the Folch method (16). Lipids were resuspended in 0.1% Triton X-100 and TAGs were determined using a colorimetric triglyceride determination kit (Sigma). True triglycerides were calculated according to a standard curve and the manufacturer’s instructions.

### Statistical analysis

All statistical testing was performed using GraphPad Prism. Statistical analysis was performed using either paired t-test, ANOVA or linear regression depending on the experimental design. Tukey’s multiple comparison test was performed with ANOVA. Data were presented as mean and standard deviation. The number of independent measurements is indicated in the figure legends.

## Results

To determine age-dependent effects on metabolism in adipose tissue, perigonadal and inguinal adipose tissue lysates from 2 mos, 7 mos and 20 mos male mice were screened by immunolabeling for key metabolic proteins (Figure 1A and 1B).

**Figure 1.**
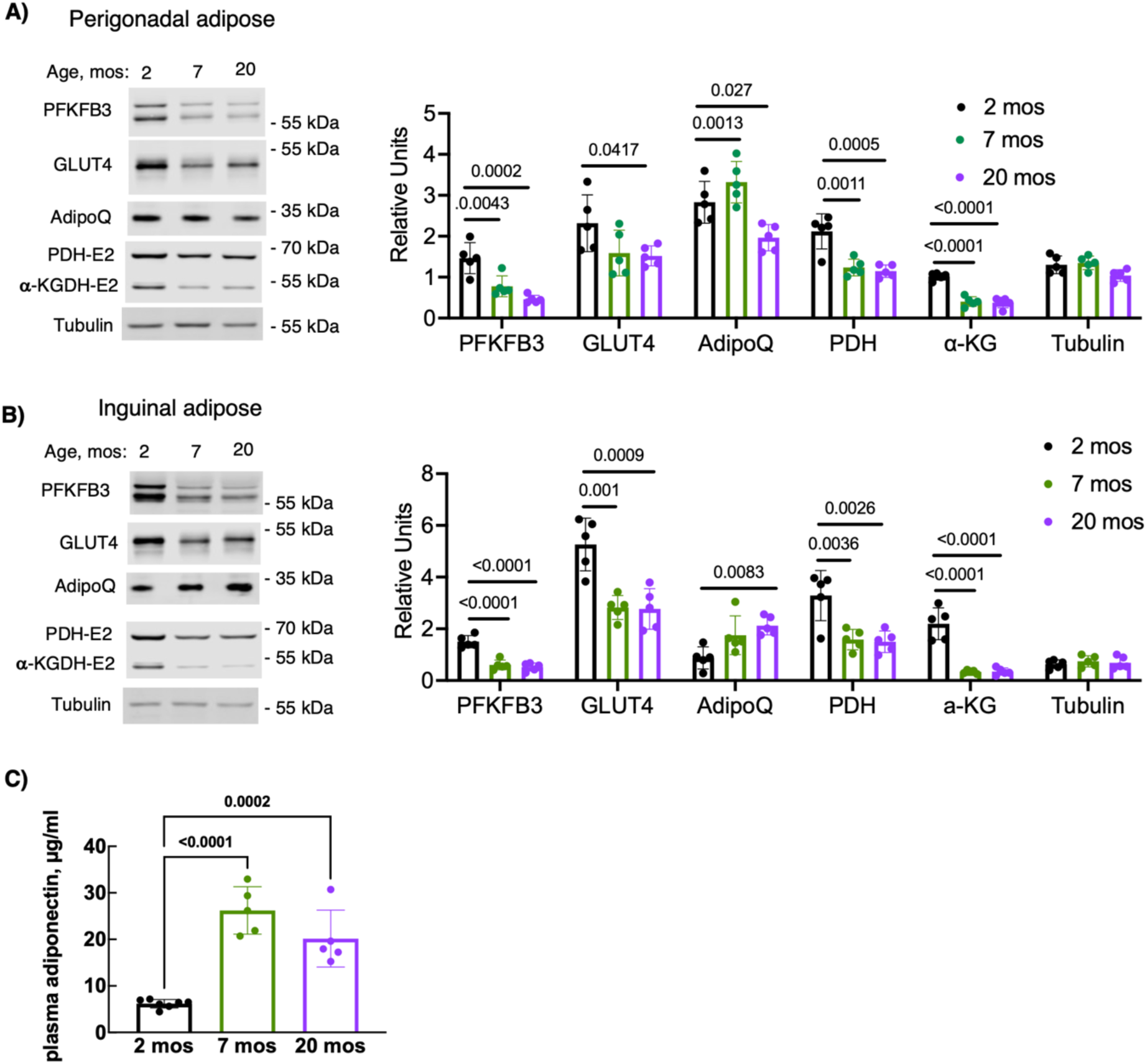
Key metabolic proteins are downregulated in aging adipose tissue. Total tissue lysates from perigonadal (A) and inguinal (B) adipose depots from male mice, age 2, 7 and 20 mos were probed for PFKFB3, GLUT4, Adiponectin (AdipoQ), the E2 subunit of pyruvate dehydrogenase (PDH-E2), the E2 subunit of alpha-ketoglutarate dehydrogenase (a-KGDH-E2) and tubulin as a loading control. Immunoblots were quantified by densitometry (n=5 per age group). Plasma adiponectin (C) was measured in fed plasma (n=5-7 per group). Data were expressed as mean and standard deviation. Data were analyzed using ANOVA with Tukey’s multiple comparisons test.

PFKFB3 protein was significantly decreased in both adipose pads by 7 mos but was not further decreased by 20 months. GLUT4 protein was also decreased as a function of age. Changes at 7 mos were more variable for GLUT4 in perigonadal tissue compared to inguinal adipose (Figure 1A and 1B). These changes predict a potential down regulation of glycolysis. To determine if mitochondrial proteins involved in glucose oxidation were similarly affected, we probed lysates for the E2-subunit of pyruvate dehydrogenase and found this protein to be decreased in the 7 mos and 20 mos adipose pads. The E2 subunit of alpha-ketoglutarate dehydrogenase was also decreased, suggesting that mitochondrial metabolism was decreased in aged fat (Figure 1A and 1B).

In contrast to PFKFB3 and GLUT4, cell-associated adiponectin (AdipoQ) levels were upregulated in inguinal adipose tissue at 7 mos and 20 mos (Figure 1). Cell-associated AdipoQ was upregulated at 7 mos in perigonadal adipose tissue, but decreased by 20 mos. We next measured plasma adiponectin in 2, 7, and 20 mos old animals and found that adiponectin levels were significantly higher in 7 and 20 mos animals compared to the very young 2 mos mice (Figure 1C). These data suggest that there is an upregulation of both adiponectin expression and secretion in aged adipose.

In a previous study, we identified PFKFB3 as a key protein regulating adipocyte function in vitro (10). PFKFB3 produces catalytic amounts of fructose 2,6 bisphosphate to accelerate the activity of *phosphofructokinase* 1, the rate-limiting step in glycolysis.

To understand the role that PFKFB3 plays in adipose tissue function and nutrient homeostasis, we generated an adipose-specific knockout of PFKFB3 in C57Bl/6 mice.

This model was generated by intercrossing mice carrying a PFKFB3 floxed allele (17) with the adipoQ-cre mouse. PFKFB3 was downregulated in the inguinal and perigonadal fat from adipocyte-specific knockout mice (Ad-KO) compared to floxed (fl/fl) controls (Figure 2A). Because PFKFB3 activity is expected to increase the rate of glycolysis, we predicted that PFKFB3 deficiency would divert glucose-6-phosphate to other pathways, such as the hexosamine biosynthetic pathway. To test this possibility, we immunolabeled periogonadal adipose lysates with an antibody against O-GlcNAc.

**Figure 2.**
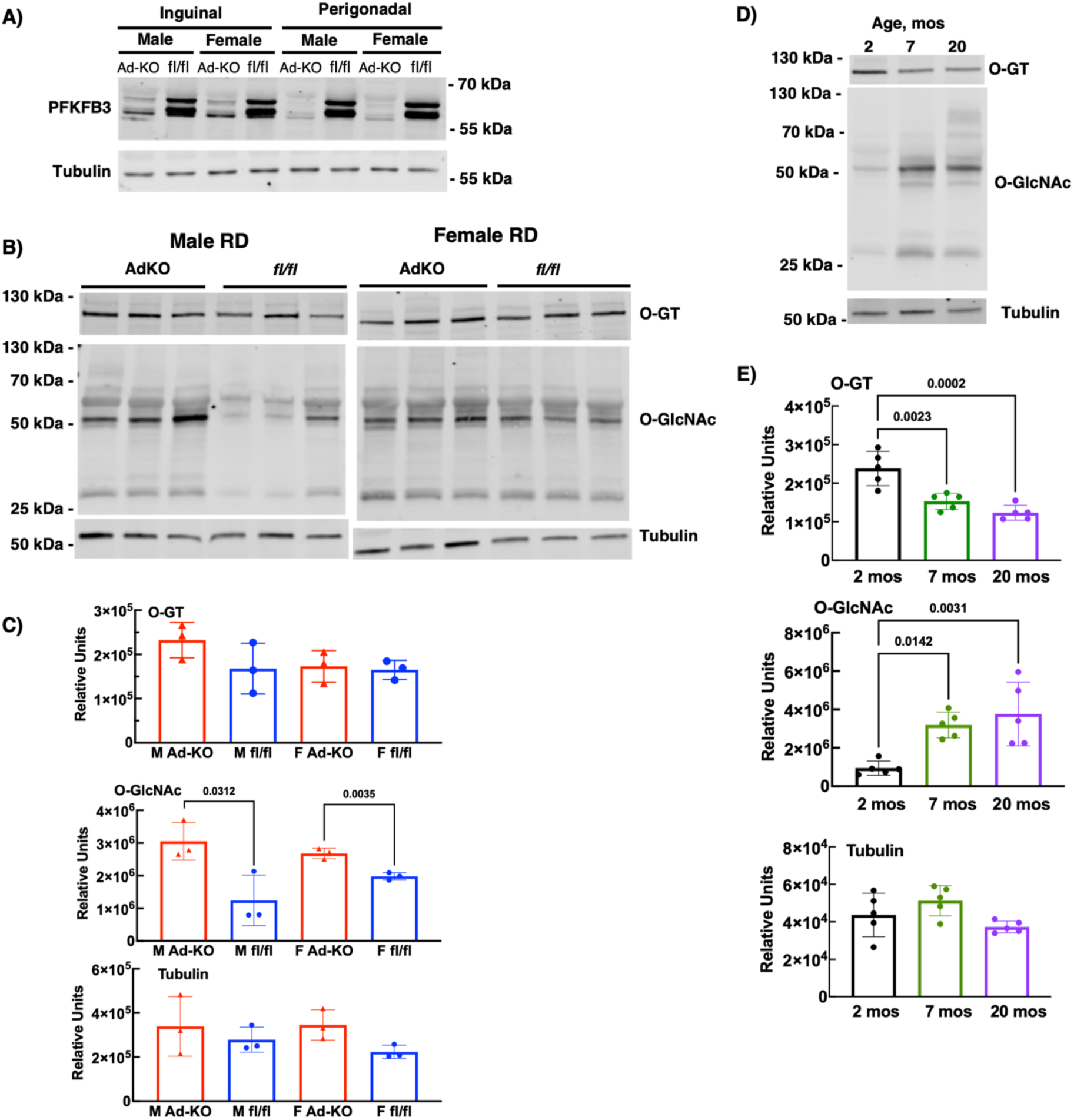
Aging and PFKFB3 deficiency in adipose tissue (A) leads to increased O-GlcNAcylated proteins (O-glcNAc). O-glcNAc, O-glcNAc transferase (O-GT), and tubulin proteins were immunoblotted (B) and quantified (C) in lysates from male and female Ad-KO and floxed (fl/fl) control mice on chow diet (RD) age 3 months (n=3 per group. O-glcNAc, O-glcNAc transferase (O-GT), and tubulin proteins were immunoblotted (D) and quantified (E) in lysates from 2-7- and 20-mo-old male mice (n=5 per group). Data are expressed as mean and standard deviation. Data were analyzed for Ad-KO mice by t-test. Data for aged mice were analyzed by ANOVA with Tukey’s multiple comparisons test.

Both RD male and female mice showed a significant upregulation of O-GlcNAcylated proteins (Figure 2B and 2C). Upregulation of O-GlcNacylated proteins in Ad-KO mice was not associated with a significant change in expression of O-GlcNAc transferase (O-GT), consistent with a change in availability of enzyme substrate rather than an increase in enzyme (Figure 2B and 2C). Like Ad-KO mice, we found that O-GlcNAcylated proteins were increased in 7 mos and 20 mos perigondal fat pads compared to 2 mos. In contrast to the Ad-KO mice, the increase in O-GlcNAcylation was accompanied with an age-dependent decrease in O-GT enzyme (Figure 2D and 2E).

To further characterize the impact of PFKFB3-deficiency in adipocytes, we assessed adipocyte area in perigonadal adipose sections from chow fed (RD) 3-mos male and female mice. Ad-KO mice of both sexes had reduced adipocyte area compared to floxed controls (Figure 3A). This suggests that lipid storage in adipocytes was decreased, limiting hypertrophic growth. When challenged with 4 weeks of 60% high fat feeding (HFD), Ad-KO male mice showed reduced adipocyte area despite an increase in overall fat mass (Figure 3B and 4A and 4E). Adipose-specific deletion of PFKFB3 did not affect plasma adiponectin levels in RD mice; however, 4-week HFD resulted in an increase in plasma adiponectin in both genotypes Figures 3C and 3D).

**Figure 3.**
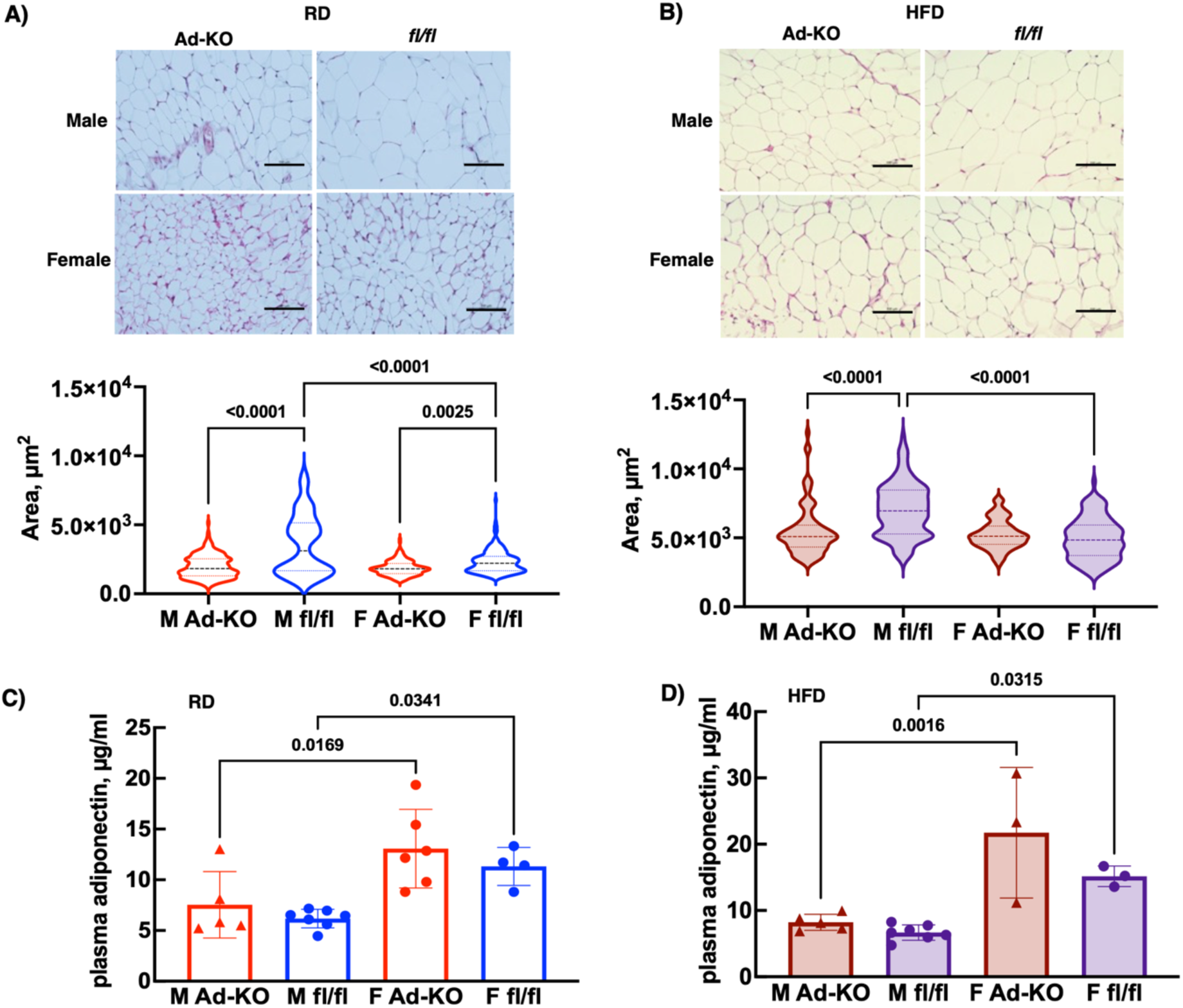
Adipose-specific PFKFB3 knockout (Ad-KO) results in decreased adipocyte area in lean (RD) 3 mos old (A) and obese (B) mice after 4 wk of high fat feeding (HFD). Area of adipocytes from hematoxylin and eosin-stained paraffin sections of male and female Ad-KO and floxed control mice quantified. Areas of 120 cells were measured by analyzing forty cells from 4 random fields each from 3 independent mice. The scale bar is 100 µM. Plasma adiponectin levels were measured in n=3-7 male and female Ad-KO and floxed control (fl/fl) mice fed Chow (RD) or 4 wks HFD. Data were analyzed by ANOVA with Tukey’s multiple comparison test. Violin plots depict the interquartile range. Adiponectin data are expressed as mean and standard deviation.

**Figure 4.**
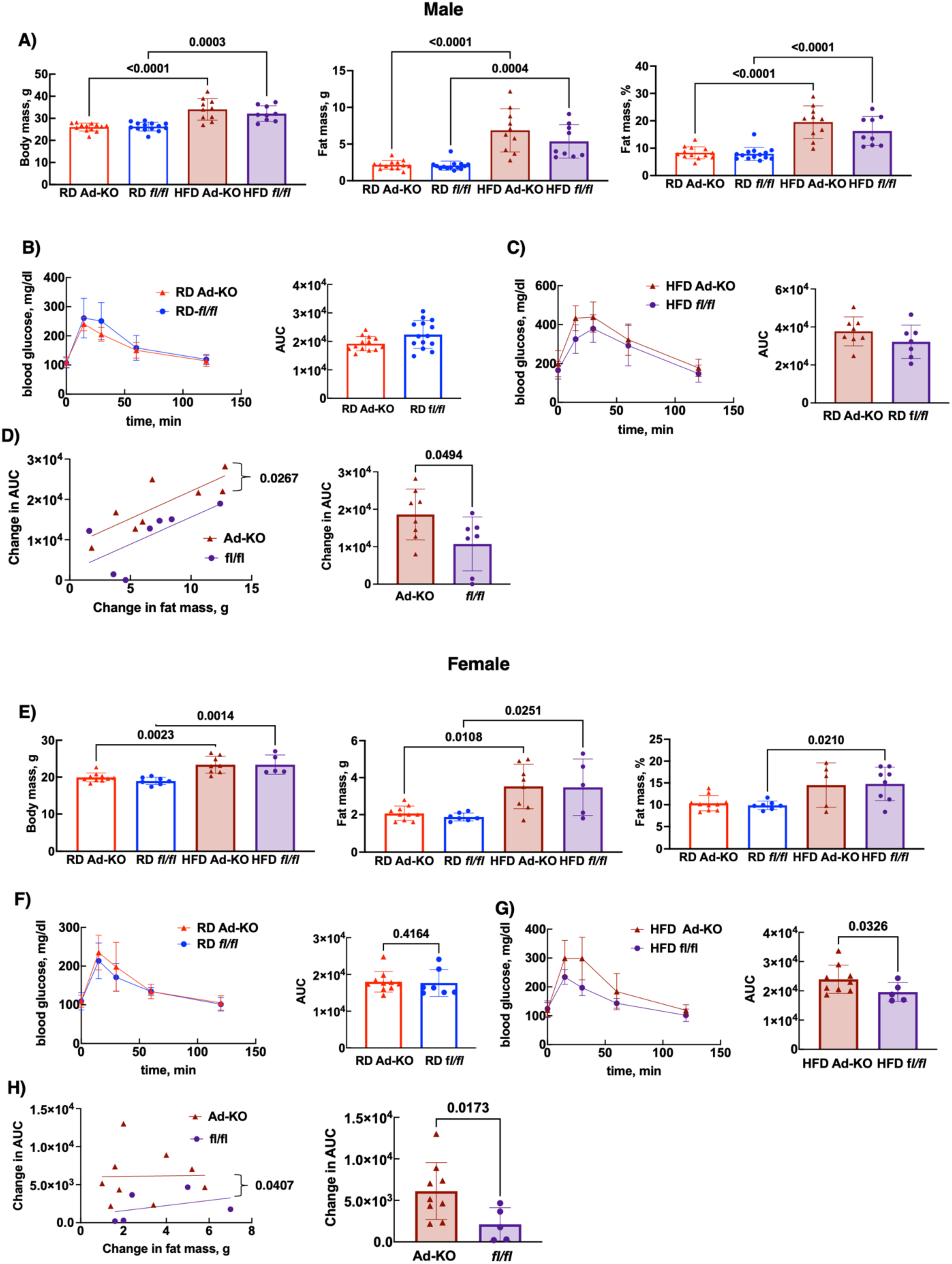
Adipose-specific PFKFB3 knockout (Ad-KO) leads to impaired glucose homeostasis with high fat diet (HFD) feeding. Body mass, total fat mass, and percent fat mass (A, E) were measured in male and female Ad-KO and floxed (fl/fl) control mice at 3 mos on chow diet (RD) and after 4 weeks on HFD. Seven to 14 mice were studied under RD conditions. Of those mice, 5 to 9 mice continued to be fed HFD for the next 4 weeks. Glucose tolerance was measured at age 3 mos (B, F), and for mice that continued with HFD, glucose tolerance was measured again 4 wks later (C, G). The change in AUC from RD to HFD for individual mice was calculated and expressed as a function of change in fat mass over the HFD period (D, H). Body composition data were analyzed by ANOVA with Tukey’s multiple range comparison. AUC for glucose tolerance and change in glucose tolerance were analyzed by t-test. Linear regression was used to analyze the change in AUC as a function of change in fat mass. Slopes of the line were similar, but the intercepts were different as indicated by the p value.

To understand how PFKFB3 deficiency impacted nutrient homeostasis, we measured body composition and glucose tolerance in 3 mo-old male and female mice before and after 4 weeks of HFD. Younger mice were used for these experiments to avoid complications associated with other age-dependent changes in other adipocyte proteins as noted in Figure 1. As expected, 4-week HFD increased body mass, fat mass and percent fat mass in both male and female mice (Figure 4A and 4E). Glucose tolerance tests were performed in each animal at 3 mos and again after four weeks of HFD. Ad-KO and floxed mice had similar measurement of the area under the curve (AUC) for the glucose tolerance test at 3 mos corresponding to the start of the study.

After the 4-week HFD, the AUC for glucose tolerance was increased in Ad-KO female mice but not in Ad-KO male. In contrast, the change in AUC over the 4-week period was significantly greater in both male and female Ad-KO mice, indicating a genotype-dependent worsening of glycemic control (Figure 4D and 4H). Because the weight gain and glucose tolerance were quite variable, we analyzed the change in in AUC as a function of the change in fat mass over the 4-week HFD period (Figure 4D and 4H). In each case the slopes of the lines were similar, but the intercepts differed (p=0.0267 for males and p=0.0407 for females) in that the Ad-KO mice had a larger change in AUC as a function of the change in fat mass.

To understand why HFD feeding worsened glycemic control in Ad-KO mice, perigonadal adipose lysates were immunoblotted for GLUT4 and PFKFB3 (Figure 5A and 5B). The RD Ad-KO female, but not male, mice had significantly lower GLUT4 protein levels. As expected, GLUT4 protein was significantly decreased by HFD in both male and female mice. PFKFB3 protein was also significantly downregulated by HFD in the fl/fl controls (Figure 5A and 5B). Because GLUT4 was down-regulated in both phenotypes under HFD, this does not explain the worsening of glucose homeostasis in the Ad-KO HFD mice. Another possible factor leading to poor glycemic control is liver triacylglycerol content (18). To test this possibility, we measured triacylglycerol concentration in livers from the RD and HFD mice (Figure 5C and 5D). While HFD did increase liver triacylglycerol concentration, there was no difference in liver triacylglycerol based on genotype.

**Figure 5.**
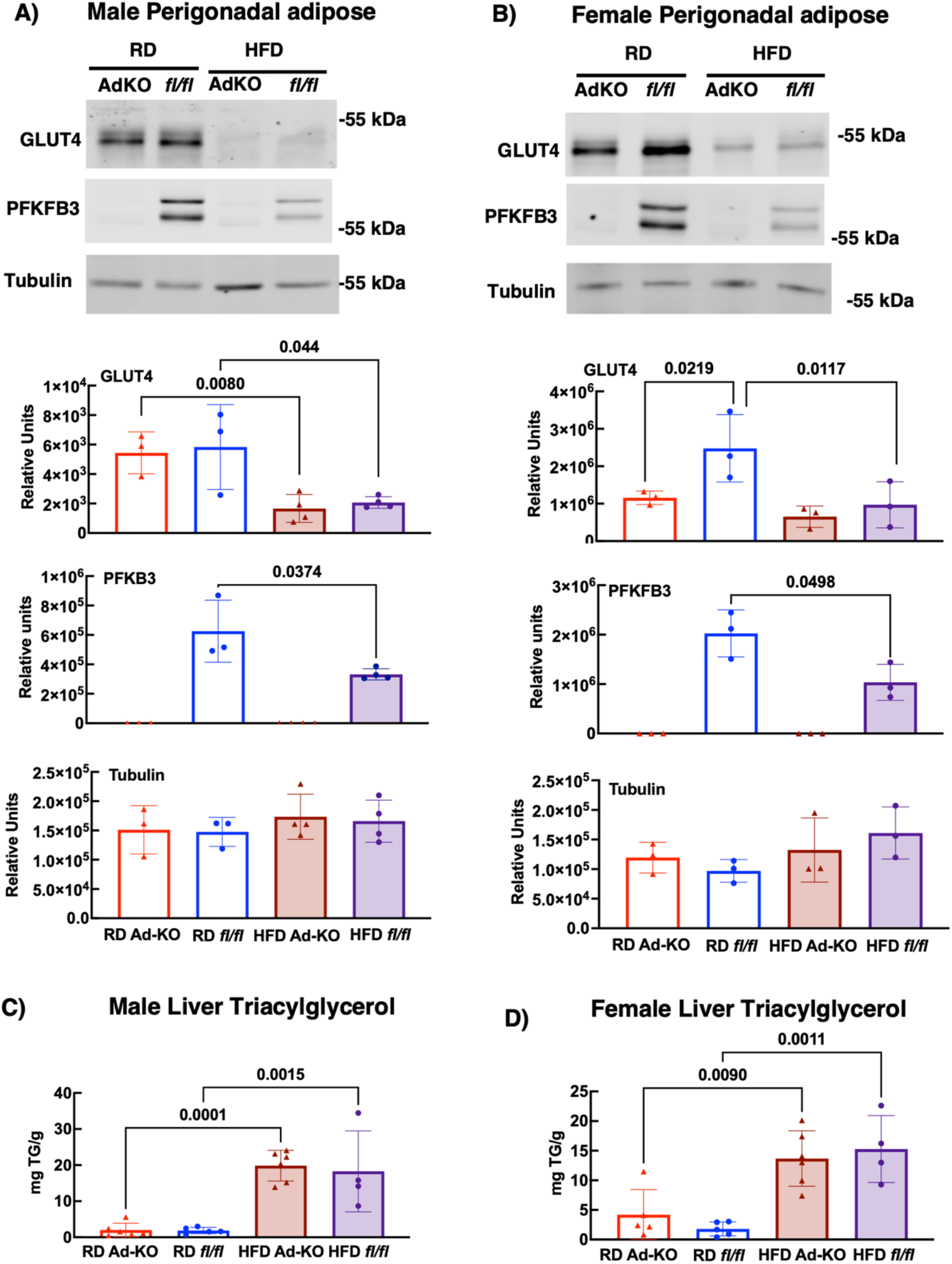
High fat feeding reduced PFKFB3 and GLUT4 protein levels in male and female mice. GLUT4, PFKFB3, and tubulin (A, B) proteins were measured in lysates from male and female Ad-KO and floxed (fl/fl) control mice on chow diet (RD) age 3 months or mice fed high fat diet (HFD) for 4 weeks (n=3 per group). Liver triglycerides were measured in RD and HFD animals (C, D). Data were expressed as mean and standard deviation and analyzed by ANOVA with Tukey’s multiple comparison test.

To test if adipose-specific PFKFB3 KO impacted adipose tissue inflammation, we Immunolabelled tissue sections of perigonadal fat ads with F480 antibody to detect macrophage infiltration (Figure 6A). F480 staining appeared to be increased in HFD tissues sections but was difficult to quantify due to the differences in adipocyte size as described above (Figure 3). To more quantitatively assess macrophage infiltration, we the blotted the adipose tissue lysates for F4/80 (Figure 6B and 6C). The tubulin loading control for these immunoblots is shown in Figure 5. F4/80 staining in both male Ad-KO RD and HFD were elevated indicating increased adipose tissue inflammation. Female mice had significantly increased F4/80 staining only in HFD conditions but not RD conditions.

**Figure 6.**
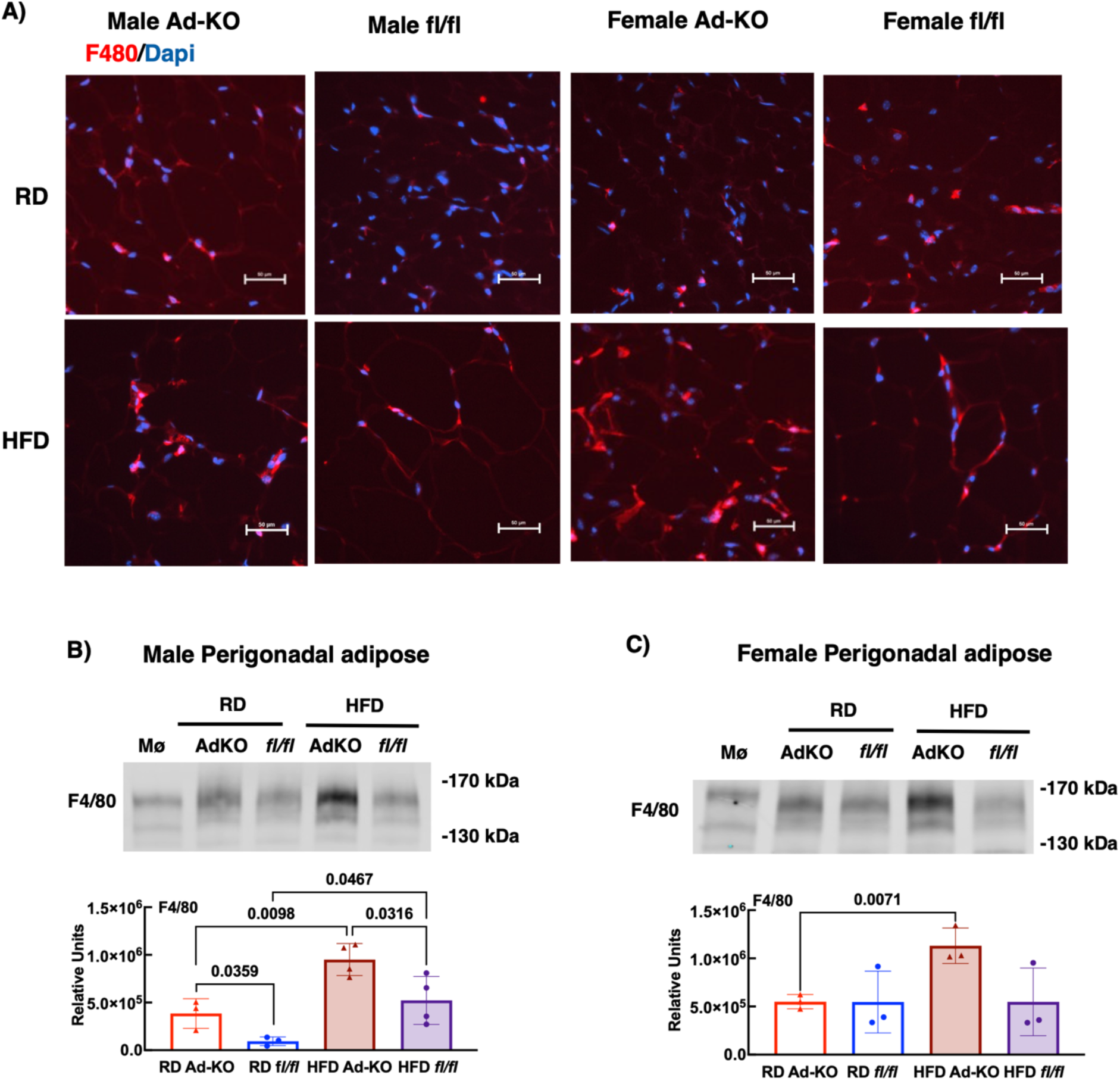
High fat diet induced macrophage infiltration was increased in adipose-specific PFKFB3 knockout (Ad-KO) mice. Selected images from perigondal adipose tissue stained with anti-F480 antibody and Dapi. Sections are from male and female Ad-KO or floxed (fl/fl) control mice at 3 mos age fed either chow diet (RD) or at 4 mos age 4 weeks of high fat diet (HFD). The scale bar is 50 µM. Images were selected form 3 random fields from 3 animals per group (A). F480 protein was measured in protein lysate from Figure 5 from male and female Ad-KO and fl/fl control mice fed RD at 3 mos or at 4 mos following 4 weeks of HFD (n=3 per group). Data were expressed as mean and standard deviation and analyzed by ANOVA with Tukey’s multiple comparison test.

## Discussion

Age-dependent changes in adipose tissue are evident by 7 mos of age in mice. Little difference is observed in expression of key metabolic proteins in white adipose tissue in 7 mos compared to 20 mos mice, two ages that represent young adult mice with a near 100% survival rate and aged mice with a near 70% survival rate, respectively (19). This suggests that changes in adipose tissue associated with aging seem to occur prior to a time when survival begins to decline. The observed changes in protein expression may represent a change in function, but not necessarily a dysfunction. For example, changes in plasma and tissue-associated adiponectin increased in the young adult and aged mice, while protein expression of several glucose metabolic genes were down-regulated.

While several changes in protein expression in aged adipose were noted, we focused on the downregulation of PFKFB3, which is a key regulator of the rate of glycolysis in in adipose tissue through production of fructose-2,6-bisphosphate (20, 21). To focus on changes that could be attributed to PFKFB3 protein deficiency and, therefore decreased glycolysis, we characterized the phenotype of adipose-specific PFKFB3 knockout mice. Importantly, we found indirect evidence that glycolysis was significantly decreased, namely that O-GlcNAcylation of proteins was increased in both the aged mice and the Ad-KO mice. This increase in O-GlcNAcylation could be attributed to increased flux of glucose-6-phosphate through the hexosamine pathway because levels of O-GT enzyme were not increased. The hexosamine pathway is an ancillary glycolytic pathway that would be expected to increase when the ratio of glucose uptake to the rate of glycolysis is increased (22). Increased hexosamine biosynthesis results in the O-GlcNAc modification of cytosolic proteins, which can change the function of those proteins. The role of O-GlcNAcylation has been probed in adipose tissue using adipose-specific in O-GT knockout mice (23). Loss of O-GlcNAcylation resulted in an inability of the fat pad to expand in response to an obesogenic diet, consistent with the role of this posttranslational modification as an intracellular nutrient sensor. The increased O-GlcNAcylation in the Ad-KO mice could explain how fat mass was not different even though fat cells were significantly smaller. It is possible that increased O-GlcNAcylated proteins play a role in signaling recruitment of preadipocytes to proceed through adipocyte differentiation.

Adipocyte area was decreased in both male and female lean Ad-KO mice. This suggested a potential change in triglyceride turnover in the Ad-KO adipocytes. Increased turnover is either due to decreased filling or increased release of triacylglycerol. We predicted that decreased filling would lead to either increased accumulation of triacylglycerol in liver or an increase in adipocyte number to accommodate the excess lipid. We favor the latter because the fat mass was not different between genotypes even though the cell size was reduced. Furthermore, liver triacylglycerol levels were not different between genotypes. When challenged with a high fat diet, the cell size in all groups increased, indicating that PFKFB3-deficient fat cell area can increase. The increase in adipocyte area was significantly smaller in Ad-KO males compared to the floxed male controls. This genotypic difference in adipocyte area was not observed in HFD female mice, likely due to the lower weight gain in the females.

The impact of adipose-specific PFKFB3 deficiency on glucose homeostasis become evident following the stress of HFD. There was no genotype-specific difference in glucose tolerance RD mice. After these mice were fed HFD for 4 wks, both the male and female Ad-KO had a greater worsening of glucose tolerance. Regardless of the level of diet-induced fat gain, the Ad-KO mice had a greater decrement in glucose tolerance. This is consistent with the notion that the PFKFB3 deficiency in adipose tissue inhibits optimal function. HFD has numerous effects on adipose tissue. We have demonstrated that GLUT4 protein expression is significantly downregulated in the HFD adipose tissue (Fig. 5A and 5B) (24, 25). In the current paper, we also report that HFD feeding reduced PFKFB3 expression in the floxed mice. The downregulation of both GLUT4 and PFKFB3 in response to a high fat diet suggest that this may be a compensatory mechanism in response to carbon overload.

This compensatory mechanism may be related to increased macrophage infiltration that occurred under HFD conditions. Importantly, worse glycemic control in HFD mice overall was correlated with increased expression of F4/80 protein, a marker of macrophages and a reduction in PFKFB3 expression. Both male and female HFD Ad-KO, which experienced even lower PFKFB3 levels, each showed a significant increase in F4/80 staining compared to the floxed counterparts, which were correlated with a further exacerbation of glycemic control at every level of adiposity. Increased adipose inflammation has previously been observed in the whole body heterozygous knockout of PFKFB3 (13). Adipose-specific overexpression of PFKFB3 suppressed inflammation, suggesting that a decrease in PFKFB3 in adipocytes is sufficient to increase adipose inflammation (14).

In conclusion, PKFKB3-mediated glycolysis impacts adipocyte biology in both lean and obese conditions. We know show PFKFB3 protein expression is down-regulated under a variety of physiologic conditions including fasting (10), aging and high fat feeding. This has an impact on adipocyte biology regarding regulation of adipocyte size, adipose inflammation and whole-body glucose homeostasis. Taken together, our data suggest that PFKFB3 in adipose tissue plays an important role in nutrient homeostasis.

## Abbreviations

AUC: area under the curve
F-2,6-BP: fructose-2,6-bisphosphate
HFD: high fat diet
O-glcNAc: O-glycnacylation
O-G: O-glcNAc transferase
PFK1: 6-phosphofructokinase-1-kinase
PFKFB3: 6-phosphofructo-2-kinase/fructose-2,6-bisphosphatase 3
RD: chow diet

## Author Contributions

BAG and ALO conceived the experiments, researched data, analyzed the data, and wrote the manuscript.

## Acknowledgements

The work was supported by a grant from the National Institutes of Health (AG079945) to ALO. The content is solely the responsibility of the authors and does not necessarily represent the official views of the National Institutes of Health. We thank Benjamin S. Harris for technical assistance.

